# Community Standards for Open Cell Migration Data

**DOI:** 10.1101/803064

**Authors:** Alejandra N. Gonzalez-Beltran, Paola Masuzzo, Christophe Ampe, Gert-Jan Bakker, Sébastien Besson, Robert H. Eibl, Peter Friedl, Matthias Gunzer, Mark Kittisopikul, Sylvia E. Le Dévédec, Simone Leo, Josh Moore, Yael Paran, Jaime Prilusky, Philippe Rocca-Serra, Philippe Roudot, Marc Schuster, Gwendolien Sergeant, Staffan Strömblad, Jason R. Swedlow, Merijn van Erp, Marleen Van Troys, Assaf Zaritsky, Susanna-Assunta Sansone, Lennart Martens

## Abstract

Cell migration research has become a high-content field. However, the quantitative information encapsulated in these complex and high-dimensional datasets is not fully exploited due to the diversity of experimental protocols and non-standardised output formats. In addition, typically the datasets are not open for reuse. Making the data open and Findable, Accessible, Interoperable, and Reusable (FAIR) will enable meta-analysis, data integration, and data mining. Standardised data formats and controlled vocabularies are essential for building a suitable infrastructure for that purpose but are not available in the cell migration domain. We here present standardisation efforts by the Cell Migration Standardisation Organization, CMSO, an open community-driven organisation to facilitate the development of standards for cell migration data. This work will foster the development of improved algorithms and tools, and enable secondary analysis of public datasets, ultimately unlocking new knowledge of the complex biological process of cell migration.

## Introduction: Towards FAIR and open cell migration data

Due to advances in molecular biology, microscopy technologies and automated image analysis, cell migration research currently produces spatially and temporally resolved, complex and large datasets. Consequently, experimental imaging techniques have *de facto* entered the “big data” era^1, 2^. This creates, on the one hand, challenges^3^ for standardising and maintaining data-driven cell migration research in public repositories while, on the other hand, offers unprecedented opportunities for data integration, data mining, and meta-analyses.

This situation resembles the progress that has been made in the *omics* fields integrating standardised data generation, sharing and analysis over the last two decades^4, 5^. The ultimate goal for cell migration data processing is to follow a similar route to progress and become more quantitative, interdisciplinary and collaborative.

To enable cell migration data integration, mining and meta-analysis, we initiated an open data exchange ecosystem for cell migration research^6^. The aim was to overcome the current fragmentation of cell migration research and facilitate data exchange, dissemination, verification, interoperability and reuse, as well as encourage data sharing^7^. This should also increase the reproducibility of experiments, enable data mining and meta-analyses, and thus satisfy the FAIR principles for Findable, Accessible, Interoperable and Reusable data^8^. Public availability of both cell migration data and metadata, and designated tools to mine these data, will facilitate the understanding of complex cell functions and their relevance for clinical use in health and disease. In addition, it will attract computational scientists to the field, producing *in silico* models allowing numerical hypotheses to be tested experimentally^9, 10^.

Establishing such an open cell migration data ecosystem needs community consensus on what content to report, what terminologies to use and what structured machine-readable formats to use in order to represent the experimental details, workflows and analysis results. A significant challenge is the inherent heterogeneity between experimental data: experiments are performed in a wide array of assays, at all levels of throughput, in diverse cellular models, maintained in various microenvironments, using multiple microscopy techniques and analysis methods. For example, a common read-out in a cell migration experiment is the measurement of the movement over time of cells and/or subcellular compartments. Other quantitative readouts include cellular morphology and its temporal dynamics^11^. However, there is no standard way to report this information, preventing the integration and mining of these data for downstream knowledge extraction. In addition, usually other experimental details are presented in narrative form in manuscripts, in line with publication policies of scientific journals, and typically not delivered in a uniform machine-readable form. While the ‘Methods’ section of scientific publications is supposed to enable full understanding of the experimental procedures and support replication, similar experiments may be described in an inconsistent manner in different studies. The methods description may be partial and may leave room for multiple interpretations of the experimental details and procedures of data analysis.

With the ultimate aim of an open data ecosystem, the Cell Migration Standardisation Organization (CMSO) was established in 2016 to define and implement standards for the cell migration community. The CMSO operates openly and transparently, is based on voluntary efforts from the community, and is open to anyone interested in contributing and/or providing feedback. The developed standards are designed and implemented aiming to achieve participants’ consensus. The CMSO outputs can be found in GitHub (https://github.com/CellMigStandOrg), while general information and activities are available from the CMSO website (http://cmso.science).

The cell migration community standards are composed of three modules, corresponding to the CMSO working groups (WGs) (**Figure 1**):

1. Reporting guidelines specifying the minimal information required when describing cell migration experiments and data (WG1);
2. Controlled Vocabularies (CVs) that unambiguously annotate these units of information (WG2);
3. Standard file formats for data and metadata, embodying the minimum reporting requirements and CV specifications, and Application Programming Interfaces (APIs), which ensure that all data, results and associated metadata can be read and interpreted by relevant software packages (WG3).

**Figure 1.**
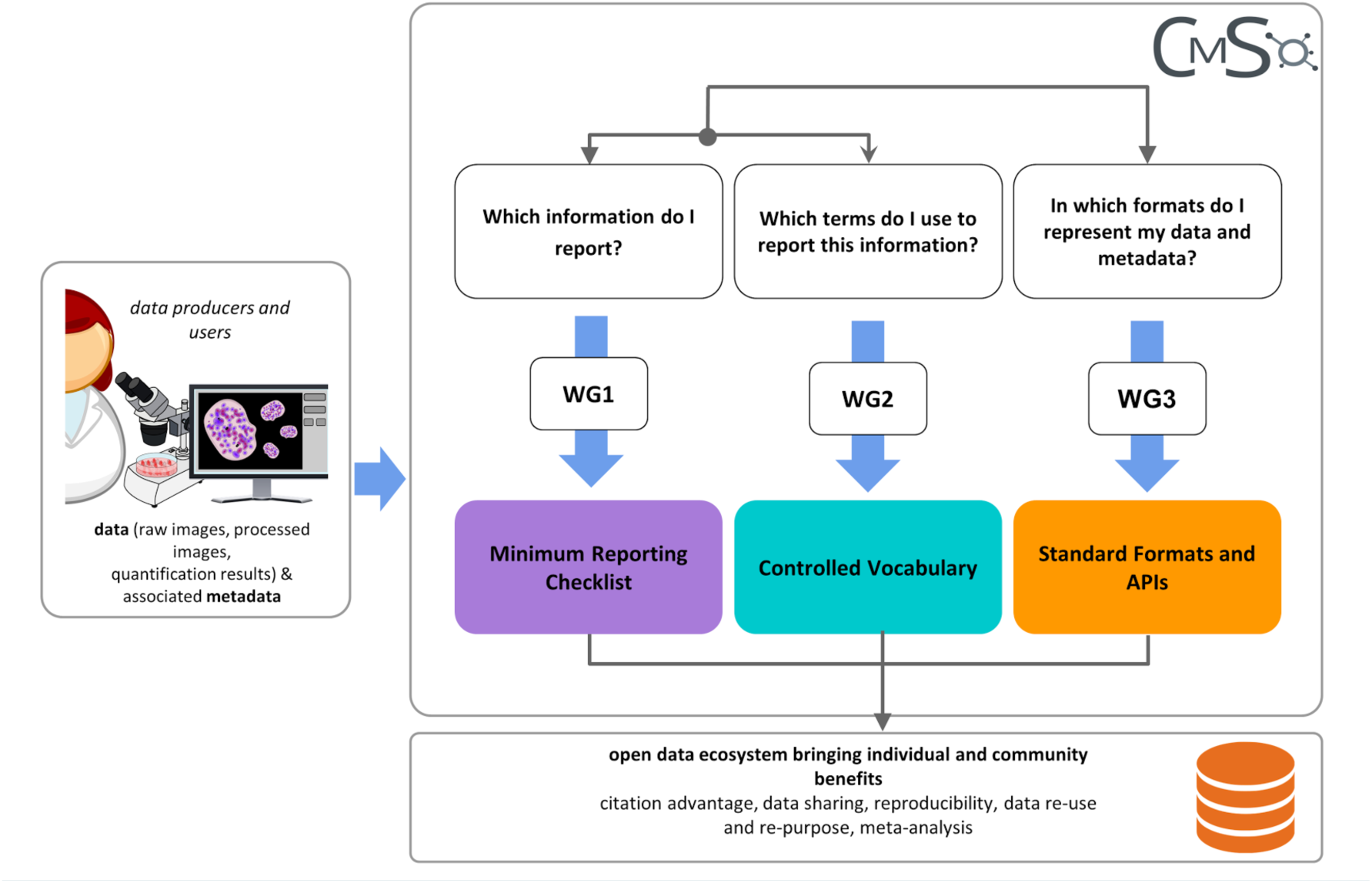
The Cell Migration Standardisation Organization, CMSO. The three working groups (WGs) deliver specific standards in an interactive manner.

While CMSO consists of these three working groups (WGs) (**Figure 1**), interactions and synergies between them were essential to achieve the integrated model. For example, the information elements identified by the minimum reporting checklist (WG1) were annotated with the terms identified as CVs (in WG2), and both checklist and vocabularies were considered when developing formats and APIs (WG3).

In this article, we introduce the CMSO standards framework, providing an audit trail from the data to the machine-readable and interoperable metadata in a harmonised manner.

## Results: CMSO standards and tools

### Minimal reporting guidelines and controlled vocabularies

Minimal reporting guidelines, also termed “requirements” or “checklists”, aim to ensure that necessary and sufficient metadata are provided to enable the comprehension of an experiment, future data integration, data mining, and to ensure reproducibility. The CMSO defined iterative versions of the Minimum Information About a Cell Migration Experiment (MIACME) guidelines, the latest version being MIACME 1.1^12^ (http://cmso.science/MIACME/, also registered^13^ in the FAIRsharing portal^14^).

MIACME consists of: (i) generic information about an investigation, which can involve one or more studies, the associated publications, people, organizations and grants, and (ii) specific information about the associated cell migration experiments. The cell migration-specific part of MIACME is partitioned into three conceptual domains (**Figure 2**): (1) the experimental setup: the assay, cell model, environmental conditions and perturbations; (2) the imaging condition: the microscopy settings; and (3) the data: the raw images, summary information about the data (e.g., number of replicates, number of images), processed images and the derived quantitative analysis outputs.

**Figure 2.**
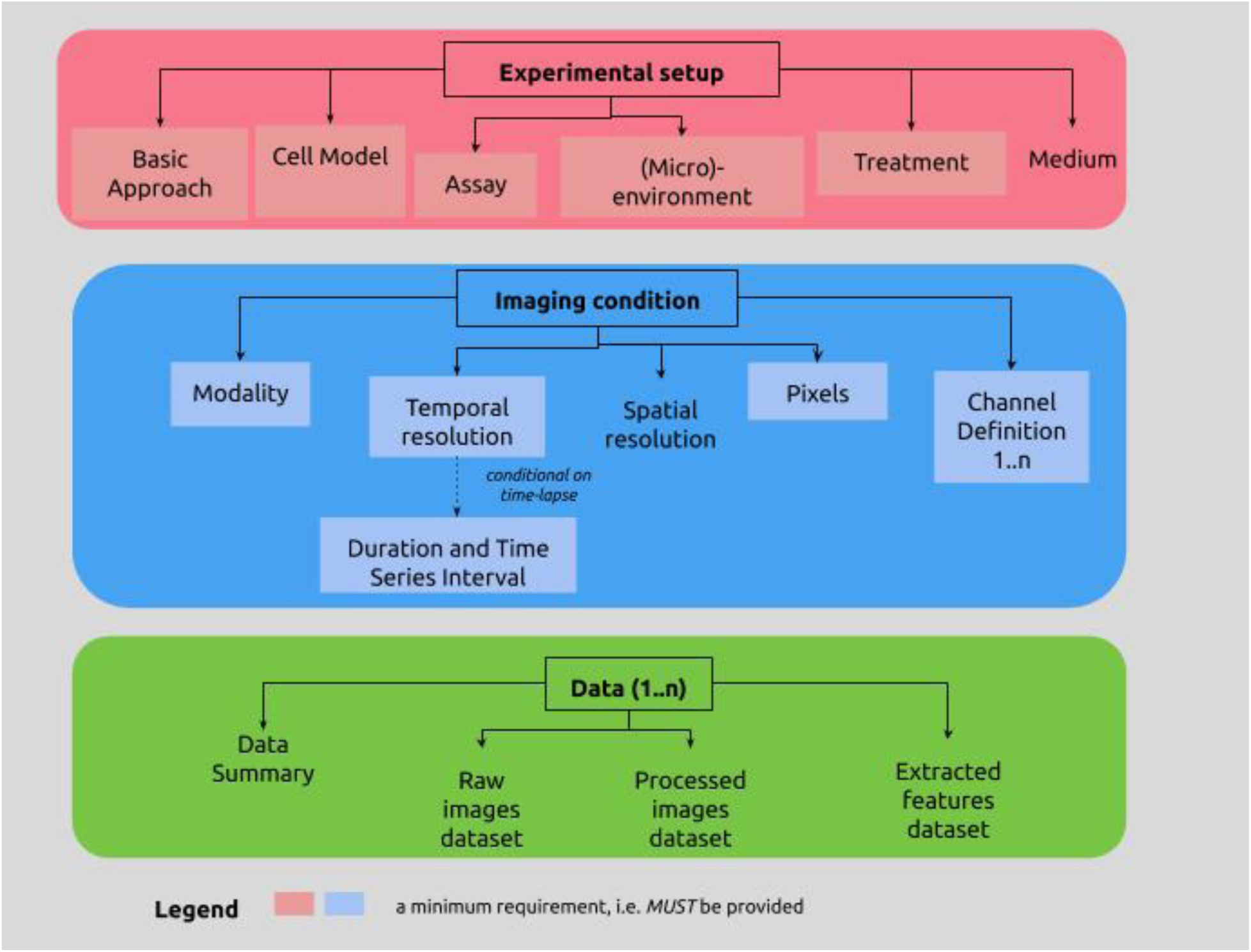
Overview of the cell migration-specific part of the MIACME specification (version 1.1^13^). The figure presents an overview of the three main components of the cell migration experiments information: experimental setup, imaging condition and data. For more details about the MIACME guidelines, including the interrelationships between the three components, see the associated spreadsheet and schemas, which specify the parameters for each conceptual area together with their requirement level, and illustrate them with examples.

MIACME is presented as a specification accompanied by a spreadsheet that describes entities and properties, their expected values, cardinalities and requirement levels. In addition, we also provide a machine-readable and actionable representation that can be validated in the form of JSON-schemas (https://json-schema.org/) and configurations (or templates) for the ISA-Tabular format, so that MIACME-compliant metadata can be created using the ISA framework^15^ (see more details in the next section).

While the minimum information requirements determine the metadata elements to be reported, the community also needs to agree on the terms that will be used when describing cell migration experiments. A controlled vocabulary (CV) provides a standard terminology with unambiguous meaning for a particular domain with the goal of promoting consistent use of terms within a community^16^. These terms can then be included in an ontology that defines formal relationships between them. CVs and ontologies harmonise the data representation to perform queries across data repositories, enable data interoperability and facilitate data integration, data mining and knowledge discovery. A typical data mining study could be based upon a set of competency questions, such as: (i) find all *in vitro* cell migration experiments that make use of live-cell imaging, (ii) retrieve all experiments for which speed was recorded for cells migrating in a 3D collagen matrix, (iii) what is the migratory effect of knockdown of gene X in a breast cancer cell line? (iv) what is the dose response of compound Y on cell line Z in an invasion experiment?

The scope of a CV is therefore defined by a list of use cases as above. For cell migration, a CV requires, for example, terms for the cell line or type, gene names, and specific compounds used for molecular intervention as well as terms describing the type of cell migration assay or the manner in which the cells are presented at the start of the assay (currently termed *cellInput* in the specification). For particularly complex experiments, additional specific terms may be needed. For instance, for single-cell chemotaxis experiments, the CV needs to include terms for the directional chemoattractant application and the type of microscopy used.

The CMSO recommends the use of multiple ontologies for reporting cell migration experiments. The selection of relevant ontologies was based on an iterative strategy, as follows.

1. Determining the domain and scope of the terminologies, through a list of possible queries used for data mining in the domain, such as those mentioned above.
2. Reusing existing terminologies. Besides being more effective, reusing terminologies is also a best practice and a requirement to interoperate with other applications that have already committed to particular CVs. The CMSO has identified existing controlled terminologies, recognized and maintained by the scientific community, which contain terms relevant for cell migration (see https://fairsharing.org/collection/CellMigrationStandardisationOrganisation).
3. Identifying missing terms (related to cell migration), specifying their definitions and relationships with existing terms. These terms are submitted to existing ontologies, when relevant, or will be created if necessary.

### Standard Formats, APIs and tools

Once the content and terminology for reporting cell migration experiments have been defined, the community needs to reach consensus on the definition of a data exchange format to enable data sharing across researchers, institutes, software tools and data repositories, as well as software libraries and APIs to interact with this format.

A single overarching file format may not suffice to capture the full complexity of the cell migration associated data, and therefore CMSO opted for a collection of well-defined, open file formats. Each format is optimized toward different aspects of the experimental pipeline: (i) experimental metadata, (ii) imaging acquisition and (iii) data analysis routines (**Figure 3**). The existing APIs and formats from the Investigation/Study/Assay (ISA)^15, 17^ and the Open Microscopy Environment (OME)^18^ are used, respectively, for the experimental metadata and the image acquisition (left and middle boxes in **Figure 3**).

**Figure 3.**
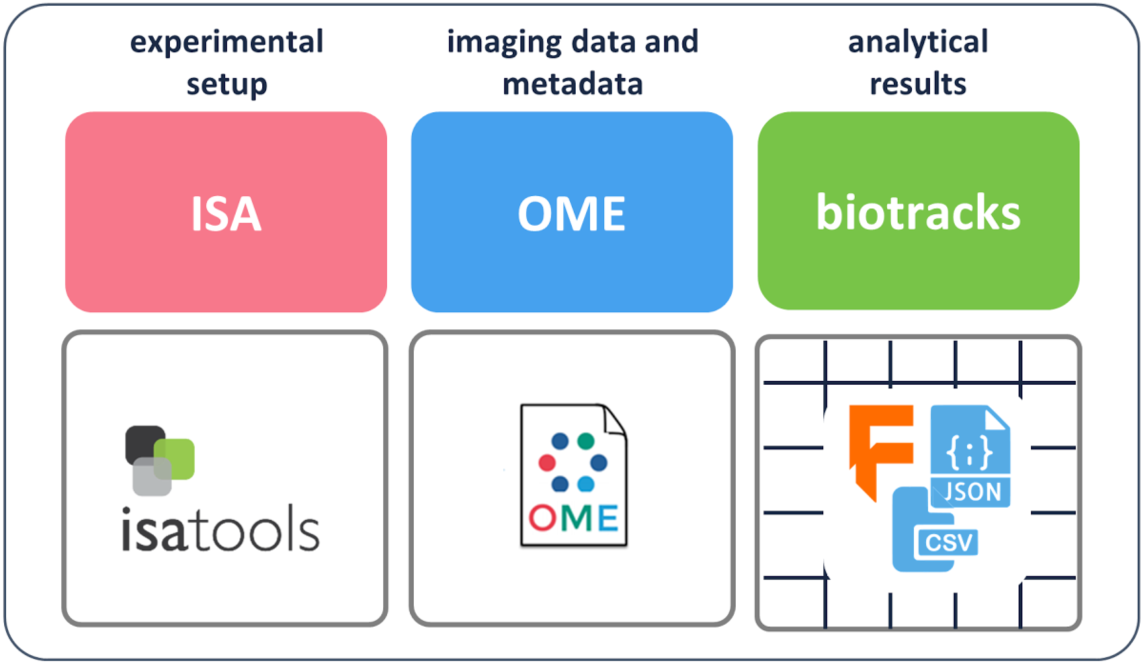
The first standardisation products assembled and developed by CMSO WG3. Experimental setup: Investigation Study Assay (**ISA)**; Image data and metadata: Open Microscopy Environment (**OME)**; Analytical results: **biotracks**.

The ISA model (http://isa-tools.org, http://isa-specs.readthedocs.org) provides a rich description of the experimental metadata (e.g., sample characteristics, technology and measurement types, sample-to-data relationships). This ISA feature served as basis for the conceptual ‘experimental setup’ section in the MIACME guidelines (**Figure 2**, pink/left box).

The OME Data Model (http://www.openmicroscopy.org/) is a specification for the exchange of image data. It represents images as 5D entities: the 2D plane (x,y), the focal position (z), the spectral channel, and the time. The OME format also includes metadata such as details of the acquisition system and experimental parameters related to acquisition. These metadata are related to the ‘imaging condition’ conceptual area within MIACME (**Figure 2**, middle box).

With the ISA and OME models established and publicly available, the remaining challenges are around standardised reporting of routine analyses, such as cell tracking or quantification of cell shape. Here, we report the specification and implementation of a new open tracking data format named **biotracks** (http://cmso.science/Tracks/, **Figures 3 and 4**). The **biotracks** format was designed to accommodate the time-resolved tracking information of various objects observed in cell migration experiments. These tracked objects can either be cells, specific organelles or cellular structures (e.g., leading cell edges, nuclei, microtubule organizing centers, filaments, single molecules, and signals that report signaling dynamics). Generally, an object could be any region of interest (ROI), including an arbitrary mask or even a single point in space. As such, the specification includes three levels of information: (i) objects identified during cell segmentation or detection tasks, (ii) links that linearly connect objects across frames of the acquired time sequence, and (iii) tracks that connect links across events such as splitting or merging (**Figure 4A**). This abstraction enables the standardised description of a wide variety of biological tracking data.

**Figure 4.**
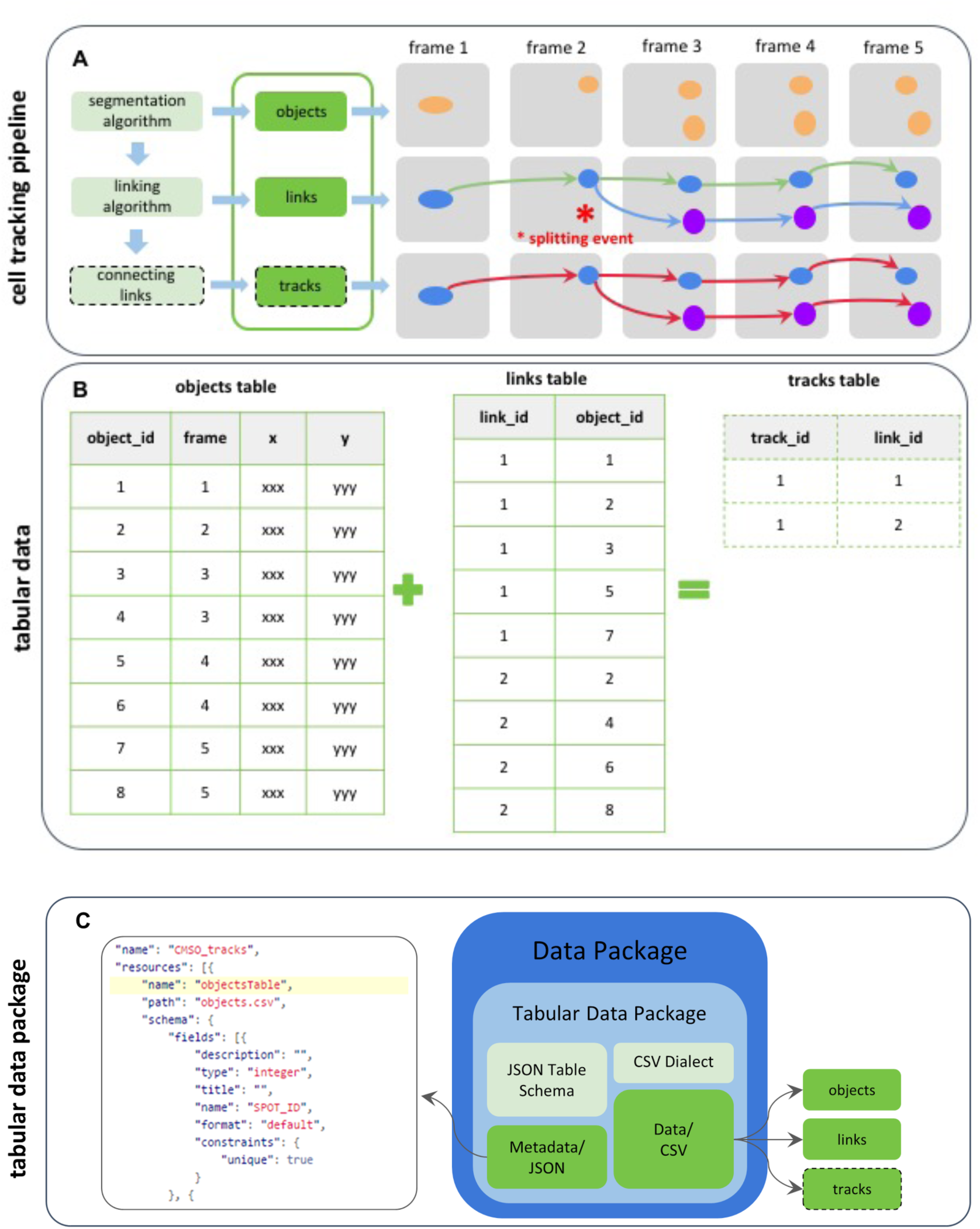
Schematic view of the biotracks format developed by the CMSO. In **A**, a segmentation algorithm identifies objects in the raw images, annotating them with the frame information, coordinates and any other features the algorithm extracts. These are described in the objects table in **B**. A linking algorithm then connects the objects across frames in a parent-child relationship. Among the possible events, the linking algorithm can then identify a split, where a parent has more than one child. This information is reported in the links table in **B**. The tracks table in **B** can finally be inferred from the objects and links tables. The tabular data package format is represented in **C**. Here, objects, links and tracks data tables are saved as comma-separated-values (CSV) files. The accompanying file in the JSON format contains both the general metadata of the data package and the metadata of the CSV files.

Whereas **biotracks** follows the general strategy of any tracking software, its specification focuses on enabling data interoperability in a simple way by specialising the Tabular Data Package (http://specs.frictionlessdata.io/tabular-data-package) container format. In the data package, the data (objects, links, and optionally tracks, **Figure 4B**) are stored in tabular form as comma-separated-values (CSV) files, while metadata and schema information are stored as a JavaScript Object Notation (JSON) file (**Figure 4C**). The development of the **biotracks** format is complementary to the OMEGA system for particle tracking data, which has particular emphasis on results from viral and vesicular trafficking experiments^19^. Standardisation is also required in other parts of the experimental process, such as standards to report how the analytical results were obtained; methods for segmentation and other image processing tasks; descriptions of post-image data exclusion and curation and descriptions of statistical analyses^20^. These aspects are left for future work by the CMSO community.

Each of the above mentioned formats (ISA, OME, and biotracks) have associated software to manipulate them, which we introduce below, together with other APIs and tools that facilitate in building machine-actionable and FAIR cell migration data.

The ISA model has an associated set of open source tools (http://isa-tools.org). In particular, the ISAcreator desktop-based tool allows for the creation, parsing and validation of experiments described with the ISA model. A version of ISAcreator is made available including MIACME configurations or templates (https://github.com/CellMigStandOrg/ISAcreator-MIACME). On the other hand, the ISA-API Python-based software (https://github.com/isa-tools/isa-api) supports the programmatic creation and manipulation of experimental metadata.

The OME model for imaging metadata is supported by several software packages, most notably the Java Bio-Formats library^21^, which can read and write OME’s own OME-TIFF standard as well as convert a wide variety of proprietary file formats into OME-TIFF (https://docs.openmicroscopy.org/bio-formats/5.9.2/supported-formats.html) and, more recently, OME-Files^22^, which serves as a reference implementation of OME-TIFF in C++ and Python.

For the analytical routines downstream, we have developed a library for cell tracking data: the **biotracks** API^23^ (https://github.com/CellMigStandOrg/biotracks). As shown in **Figure 5**, the library takes as input a cell tracking file from a tracking software (such as TrackMate^24^, CellProfiler^25^, Icy^26^ and MosaicSuite^27^) and produces a data package where objects and links are stored in the standardised format depicted in **Figure 4**.

**Figure 5.**
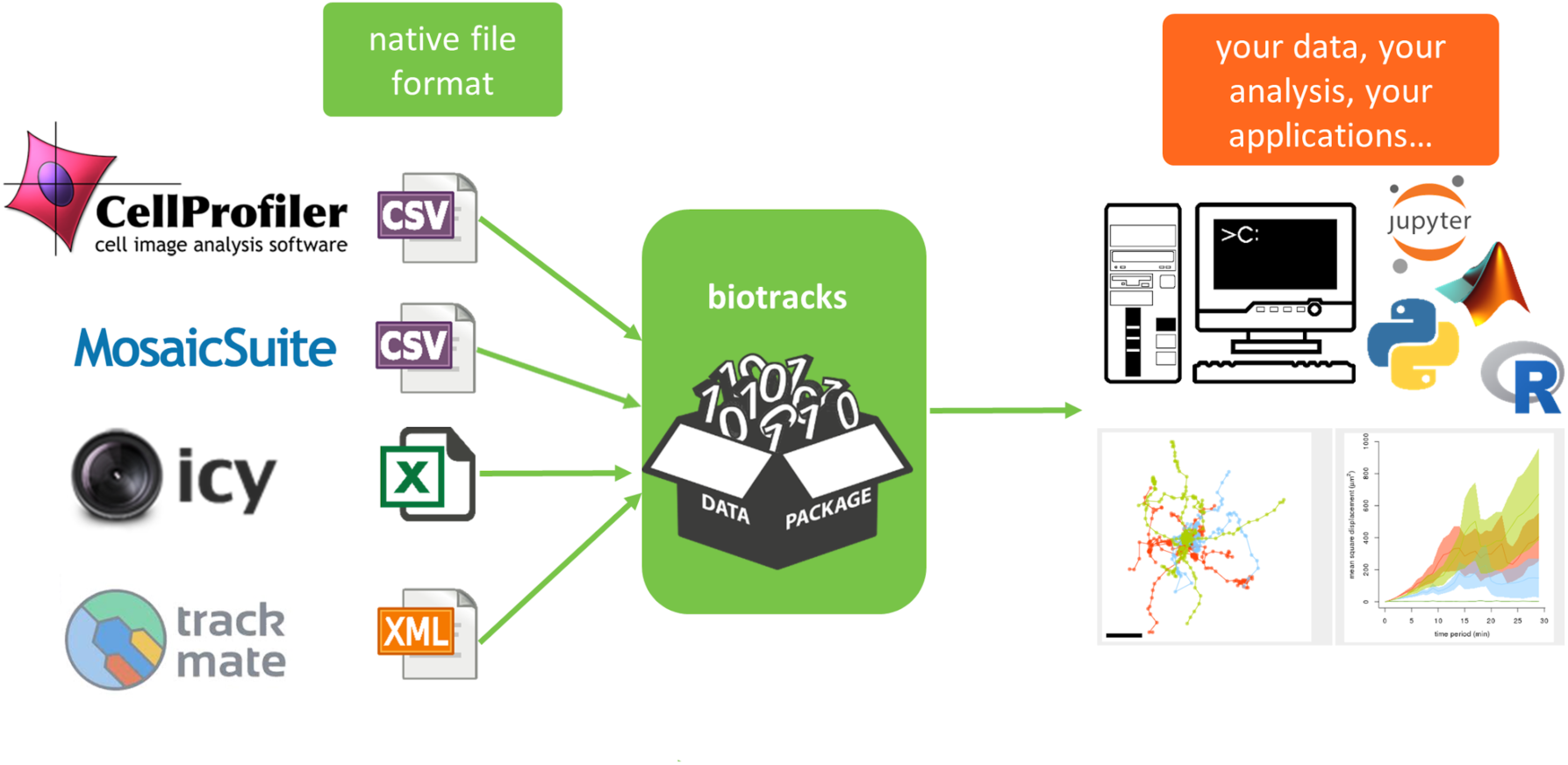
The biotracks library. The library receives cell tracking data as input from multiple tracking software and converts them to the **biotracks** format (see Figure 4), which can be further visualized and analyzed with downstream applications within this framework.

The CellMissy software package^28, 29^ (https://github.com/compomics/cellmissy), a cross-platform data management and analysis system for cell migration/invasion data, was extended to import and export datasets whose experimental metadata is available in MIACME-compliant ISA-Tab format and whose cell tracking data are represented with **biotracks**.

A cell migration data repository (https://repo.cellmigration.org/) was created, which accepts submissions of experimental metadata in ISA-Tab format compliant with the MIACME guidelines together with raw data submission, and supports searching across the deposited data.

The CMSO standards were incorporated into the WiSoft platform, a commercial software developed and distributed by IDEA Bio-Medical LTD, which consists of two software tools – Athena and Minerva - designed to support the addition of experimental parameters, imaging properties, and analysis modules. This extensibility feature facilitated the adoption of the MIACME elements and allows easy adaptation of algorithms contributed by researchers as part of the ongoing effort of analyzing dynamic biological data.

### CMSO Standards in Action

To demonstrate the application of the CMSO standards we applied them to the study by Masuzzo *et al.*^28^, which proposed an end-to-end software solution for the visualization and analysis of high-throughput single-cell migration experiments. The authors used two datasets to demonstrate their software. Here we reuse their Ba/F3 cells experiment to demonstrate CMSO standards in action as follows. As a first step, we annotated the data using the MIACME guidelines (v1.1) (see example in **Table 1** in the Supplementary Information). The two columns of this table represent the two layers of information of **Figure 2**: specific entities (column 1) are annotated using (controlled) terms (column 2). As shown through this example, the MIACME schema converts a large imaging-based study into an easy-to-interpret structured description. The resulting metadata are available in Github (https://github.com/CellMigStandOrg/CMSO-datasets/tree/master/cmsodataset0001-masuzzo).

As shown above, when reporting an experiment, researchers will need to complete the information indicated in MIACME, and for those fields that require a controlled vocabulary or ontology term, they will need to select the most appropriate terms, considering the suggested ontologies for each field (e.g., for organisms, MIACME recommends the NCBI Taxonomy). The criterion for selecting a term is to consider the most specific description available in the ontology. If no term exists in current ontologies (shown as [*] in **Table 1** in the Supplementary Information), the CMSO community has been submitting it to a relevant ontology. If there is no relevant ontology, the CMSO community has been gathering the terms to create a specific ontology in the future, if necessary.

This MIACME information is included in an ISA-Tab representation of the dataset, which also expands it with more information about the processes performed in the experiment and their intermediate inputs and outputs.

Secondly, to demonstrate the usage of the data formats and APIs, we have prepared an interactive Jupyter notebook, available at https://github.com/CellMigStandOrg/CMSO-training/blob/master/notebooks/CMSO_PM.ipynb. The notebook uses the set of three software libraries discussed above: ISA-tools to manipulate the experimental setup information, OME-files for the imaging data and metadata, and **biotracks** for the cell tracking data. The pipeline presented in the notebook bundles the three libraries together, showing the interaction of the CMSO standards in a complete experimental and analytical workflow.

## Discussion: The overall vision

Genomics, proteomics and structural biology have greatly benefited from well-developed data standards^30^ that contributed to rapid progress in these fields. However, in the field of cell migration, the lack of unifying standards and repositories have limited the opportunities to make similar progress. Thus, expensive and difficult-to-generate imaging data are stored at local labs with no further access for the community and with no standardised descriptions of the experiments that generated them. CMSO has taken initiatives to increase accessibility and reproducibility of cell migration data across models, by developing an open access reporting structure that aims to accommodate diverse types and complexities of cell migration data. We consider this as a first step towards standardisation of cell migration data, which will facilitate integration, validation and meta-analysis of cell migration data across models, and foster progress across study and model comparison, enabling the validation of new discoveries.

In this work, we presented a framework around community-driven standards and tools, developed by CMSO through an open process, for managing cell migration data along its data life cycle. The CMSO framework relies on established standards for experimental metadata (ISA) and imaging data (OME), both complemented with models and tools developed by CMSO. We introduced reporting guidelines that identify what elements should be reported for cell migration experiments (MIACME), and a format for cell tracking data. We also provide APIs and software tools supporting the description and publication of cell migration experiments, their workflows and results.

Open cell migration data following data standards and the tools to manipulate them will enhance the performance and relevance of the field and deepen insight into this complex biological progress beyond the impact of primary research. This requires future actions and tools moving forward: the community must improve the user-friendliness of the routine processes of data curation, deposition and exchange. It also needs ensuring that the data standards continuously evolve to meet community needs. Eventually, the community must strive toward making maximal use of the data: contributing public data; developing new data-driven computational tools; mining for patterns that can drive new biological hypotheses; testing new hypotheses in experimental, computational, and clinical models; and unlocking new knowledge that drives scientific progress and yields new therapies and strategies to improve health.

### Outlook: CMSO sustainability

Proper standards and broad community support are crucial for the establishment of a long term open data sharing ecosystem for cell migration research. Therefore, the output of CMSO is a crucial cornerstone for the implementation of such an ecosystem. The European Union Horizon 2020 MULTIMOT project (https://multimot.org/) has been the initial driving force for the development of CMSO, from establishing the organization to arranging the first meetings, enabling the involvement of the broader cell migration community. From the start CMSO was planned as an independent entity, including people beyond MULTIMOT, with its own governance structure (https://cmso.science/roles-and-responsibilities/) and a community-driven decisions mechanism. MULTIMOT members are required to implement the CMSO standards, and therefore they constitute the first users and quality assurance testers. CMSO members are responsible for the dissemination of the materials produced, and for the sustainability of the organization in the future through funding.

One of the prime goals of CMSO is to raise community awareness on best practices for data stewardship, to promote and disseminate the use of community standards. CMSO is managed and run by volunteers from the community, and is open to participation from anyone interested. Current CMSO participants include cell biologists, immunologists, cancer researchers, medical professionals from laboratory medicine, microscopists, computational biologists, and data scientists (https://cmso.science/how-to-get-involved/). The CMSO also welcomes cell migration data contributions from the scientific community and provides guidelines for creating MIACME-compliant descriptions of experiments and using CVs to annotate them.

## Methods

### Compilation of use cases

In order to provide incentives for cell migration researchers to invest the time and effort required to structure their data and make it FAIR, CMSO identified a series of use cases where applying the CMSO framework would enable data integration and data reuse and would drive further scientific discoveries.

Combining harmonised data (e.g. from the same cell culture model retrieved from different studies^31–34^) will facilitate data analysis and mining across imaging acquisition techniques, set-ups and cell lines, with applications such as: (i) comparison of results from 2D and 3D culture environments that make use of the same cell model; (ii) validation of *in vitro* results from against datasets from *in vivo* experiments in model organisms (e.g. zebrafish embryos, mouse models of disease) to ascertain *in vivo* relevance; or (iii) systematic comparison of the relative dose-effects of growth factors or chemical compounds on migration behaviors across cell models.

As an important further opportunity, the reuse of existing primary image data with new analyses can reveal previously unexplored patterns contained in such complex data^31–34^. Typical examples for secondary reuse^7^ of multiparametric imaging datasets results from applying novel computational algorithms to derive kinetic shape features (e.g. leading edge oscillations or membrane curvature changes) or functional components such as switch behaviours and stochasticity in cell population behaviour^9, 35^. Furthermore, public datasets can serve as benchmarks for comparing the performance of computational and analytical methods. When they include proper metadata annotation, such datasets are invaluable for developing new methods^36–38^ and training machine learning algorithms in a variety of analytical tasks ^39^. As resources for the development and comparison of methods, the value of large amounts of publicly available image data cannot be overstated^40–42^.

### FAIR standards for cell migration

Since the turn of the century, there have been standardisation efforts in different domains: genomics^43, 44^, proteomics^45–47^, and metabolomics^48–50^. Traditionally, these efforts considered each of the aspects (content, terminology, formats) independently, see the FAIRsharing repository of standards^30^. Different communities developed minimum information guidelines in narrative form (e.g., MIAME for microarray^51^; MIAPE for proteomics^45^) and while they encouraged the use of ontologies^52^ and formats^46^ for annotation and representation, respectively, they developed them separately and emphasised that the guidelines were implementation-independent^45^. Whereas independent reporting guidelines imply that they can be implemented in different formats with different semantic models, the importance of producing data that is truly FAIR in cell migration and the breadth of assays and technologies that are used in the field mandates that a reporting guideline in narrative form is no longer sufficient. The need for FAIR data requires a reference implementation, i.e. a standard software tool(s) that reads and writes in a standardised format that meets the specification, along with clear, usable examples that show how to implement and use the format. Finally, it is essential to provide validation tool(s) so that scientists and technology developers attempting to adopt the standard can be assured their work is compliant with the reporting guidelines. A comprehensive approach that encompasses the content, terminology and format and thus considers the full machine-actionable model for data description is needed. This is the strategy chosen by CMSO. In this way, the minimal reporting checklist is a first step towards the identification of the metadata elements required for a full data description model, especially in view of the FAIR data principles^8^ and FAIR data models^53, 54^. We chose a variety of formats to represent the checklist, including a machine-actionable and FAIR representation based on JSON-schemas for JSON-LD data.

To produce these models and tools following a community-driven approach, during face-to-face workshops as well as via online tools, we run the following activities:

- Identified metadata descriptors for cell migration experiments based on the model used by the CellMissy software tool. These metadata descriptors were divided into three categories: experimental setup, imaging condition and cell migration data. Then, a group of cell migration researchers ranked the descriptors according to three values representing: important, somewhat important and not so important. The analysis of the ranking provided the first guidance to build the initial MIACME guidelines.
- After choosing a representative paper on cell migration^55^, a survey was prepared and distributed to researchers asking them to complete the values of the identified metadata descriptors considering the experimental description given in the paper. By asking them to complete the values, we could check if it was easy to identify the relevant element in the paper, and how clear the explanation about the descriptor was. The descriptors were split in multiple sections: general experiment overview and description, cell system description, cell culture conditions, assay description, vessel, plate and environment information, perturbation and intervention, imaging, image analysis information, licensing and terms of use, and request for feedback. Again, we requested researchers to rate each element on a 1 to 5 scale, from essential to useless.
- The above steps and discussions with researchers allowed us to refine the metadata descriptors to be included in MIACME, and in this way we developed several versions of the checklist.
- During this iterative process, we also identified the values for the different elements for a variety of experiments, some published and some unpublished. The feedback and discussions among researchers about the importance of each descriptor was crucial to keep refining the checklist and settle on a set of descriptors deemed minimal but also sufficient to enable the comprehension and replicability of the experiment.
- In terms of semantic annotations, we found terms in existing ontologies when available and otherwise requested the addition of terms in relevant ontologies (e.g. https://github.com/information-artifact-ontology/IAO/issues/212)
- The development of a common standard format to represent cell tracking data associated with cell migration experiments aimed to produce a simple and extensible format, reuse existing standard formats where possible and support both human- and machine-readable metadata. The adopted solution relies on the Tabular Data Package, which supports the association of data with a JSON file to specify metadata and schema information, and resulted in a specification (https://cmso.science/Tracks/) and a python-based software tool (biotracks) to manage the new format.
- We also compiled experimental datasets to demonstrate the application of the different standards and how they integrate in the CMSO framework.

## Acknowledgments

The Cell Migration Standardisation Organisation acknowledges funding from the European Union’s Horizon 2020 Programme under the MultiMot project, Grant Agreement 634107 (PHC32-2014).

## Author Contributions

AGB and PM contributed equally to writing the manuscript and addressed all authors comments and contributions. AGB led the development of the MIACME reporting guidelines, coordinating the community feedback and updates, with contributions from PRS, PM, MVT, AZ, RHE, LM and feedback from cell migration researchers involved in CMSO via face-to-face meetings and online communications. AGB wrote the MIACME specification, PRS and AGB produced MIACME-compliant ISA-Tab configurations. AGB produced MIACME JSON-schemas and JSON-LD context files. PM and SB led the development of the biotracks specification with contributions from SL, GS, JM and the CMSO community. PM led the development of the biotracks package with contributions from SL and GS. GS led the extension of CellMissy to support the CMSO standards. JP developed the cell migration repository. YP extended the IDEA Bio-Medical LTD software to support the CMSO standards. PRS produced the ISA-TAB representation of cell migration studies published by PM. PRS provided the list of recommended ontologies. PM, AGB, SL produced the examples of CMSO datasets with their MIACME, ISA-Tab, OME and biotracks representations and code to validate them. SB, AGB, SL, PM, MVT produced the training material. AGB created the FAIRsharing collection for CMSO and included its widget in the CMSO website. AGB, SB, LM and PM produced the CMSO website. All authors contributed to, read and approved the final manuscript.

## Abbreviations

|  |  |
| --- | --- |
| API | Application Programming Interface |
| CBO | Cell Behaviour Ontology |
| chEBI | Chemical Entities of Biological Interest |
| CLO | Cell Line Ontology |
| CMPO | Cellular Microscopy Phenotype Ontology |
| CMSO | Cell Migration Standardisation Organization |
| CSV | Comma Separated Values |
| CV | Controlled Vocabulary |
| EFO | Experimental Factor Ontology |
| ERO | eagle-i Research Resource Ontology |
| FAIR | Findable, Accessible, Interoperable and Reusable |
| GO | Gene Ontology |
| HPO | Human Phenotype Ontology |
| ISA | Investigation, Study, Assay |
| JSON | JavaScript Object Notation |
| JSON-LD | JavaScript Object Notation for Linked Data |
| JSON-schema | JavaScript Object Notation Schema |
| MIACME | Minimum Information for Cell Migration Experiments |
| MIAME | Minimum Information About a Microarray Experiment |
| MIAPE | Minimum Information About a Proteomics Experiment |
| NCBITaxon | NCBI Taxonomy |
| OBI | Ontology for Biomedical Investigation |
| OME | Open Microscopy Environment |
| OME-TIFF | Open Microscopy Environment Tagged Image File Format |
| ROI | Region of Interest |
| UO | Units Ontology |
| WG | Working Group |

## Supplementary Information

**Table 1:**
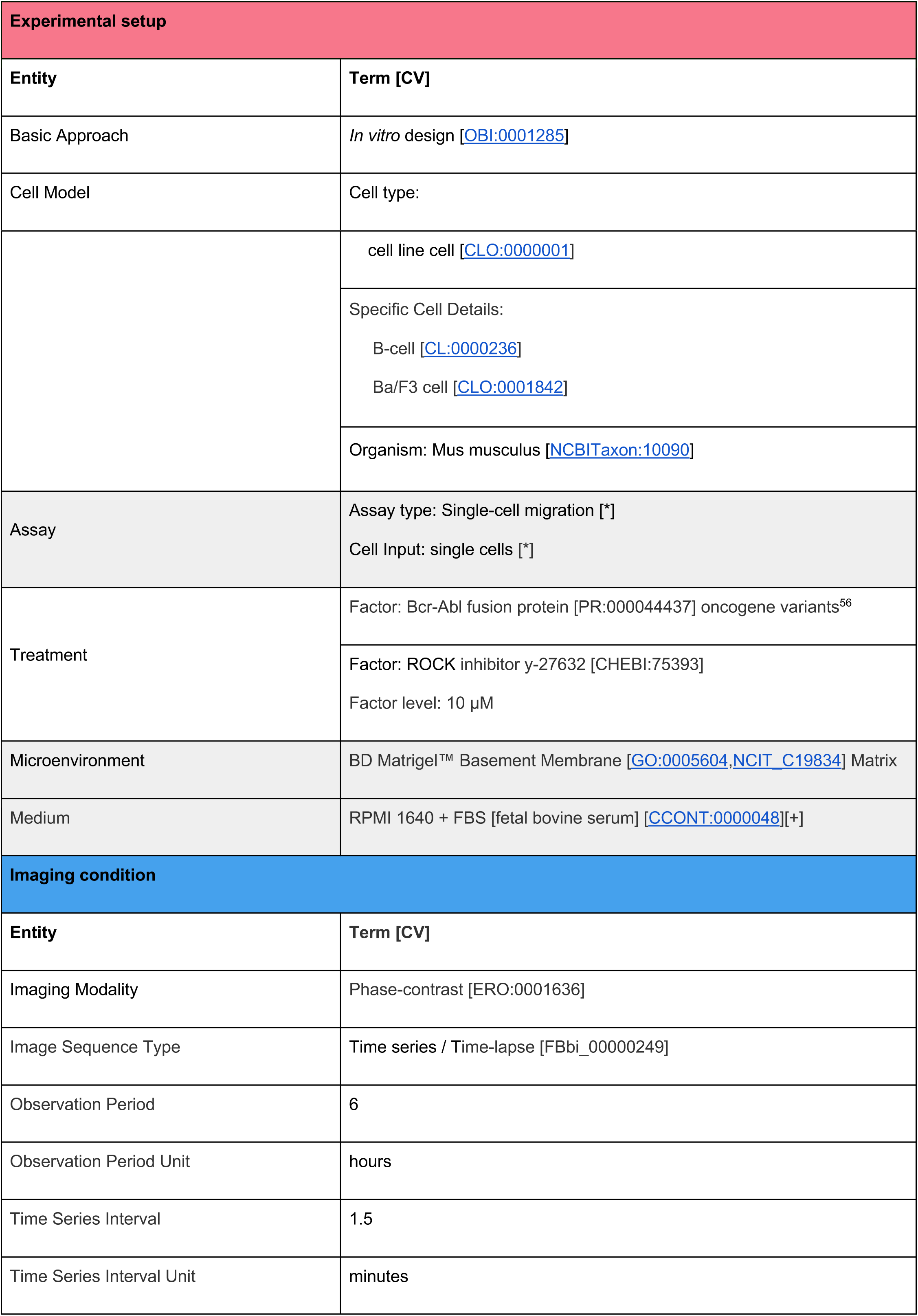
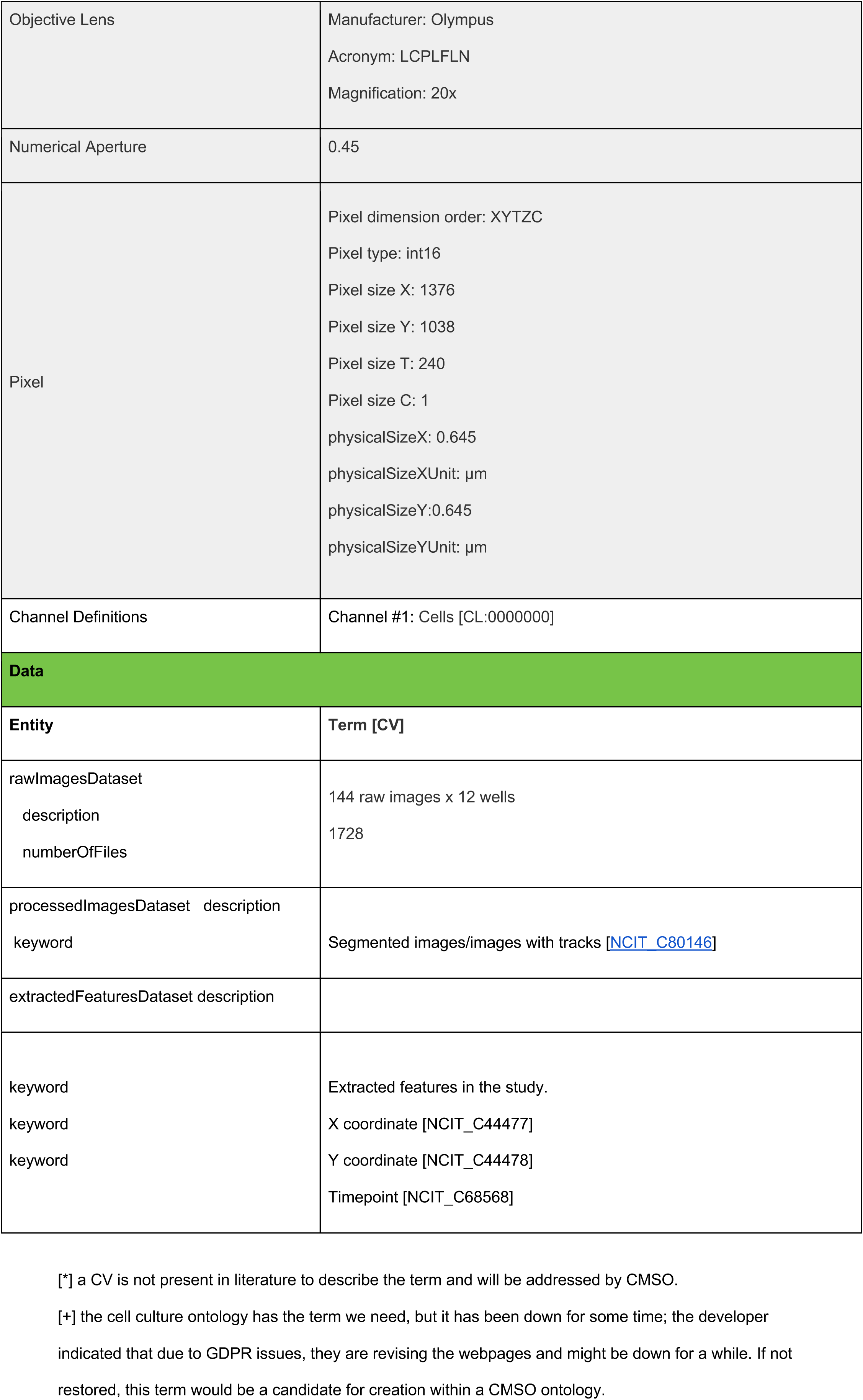
The cell migration study from Masuzzo *et al*.^28^ described and annotated using the cell-migration-specific part of the MIACME guidelines (version 1.1). Highlighted in gray, we show elements that are part of the MIACME requirements but were not reported in the original paper.

